# Probing the *in-situ* volumes of Arabidopsis leaf plastids using 3D confocal and scanning electron microscopy

**DOI:** 10.1101/2023.07.16.549198

**Authors:** Jan Knoblauch, Rainer Waadt, Asaph B. Cousins, Hans-Henning Kunz

## Abstract

Leaf plastids harbor a plethora of biochemical reactions including photosynthesis, one of the most important metabolic pathways on earth. Scientists are eager to unveil the physiological processes within the organelle but also their interconnection with the rest of the plant cell. An increasingly important feature of this venture is to use experimental data in the design of metabolic models. A remaining obstacle has been the limited in situ volume information of plastids and other cell organelles. To fill this gap for chloroplasts, we established three microscopy protocols delivering *in situ* volumes based on: 1) chlorophyll fluorescence emerging from the thylakoid membrane, 2) a CFP marker embedded in the envelope, and 3) calculations from serial block-face scanning electron microscopy (SBFSEM). The obtained data were corroborated by comparing wild-type data with two mutant lines affected in the plastid division machinery known to produce small and large mesophyll chloroplasts, respectively. Furthermore, we also determined the volume of the much smaller guard cell plastids. Interestingly, their volume is not governed by the same components of the division machinery which defines mesophyll plastid size. Based on our three approaches the average volume of a mature Col-0 wild-type mesophyll chloroplasts is 93 µm^3^. Wild-type guard cell plastids are approximately 18 µm^3^. Lastly, our comparative analysis shows that the chlorophyll fluorescence analysis can accurately determine chloroplast volumes, providing an important tool to research groups without access to transgenic marker lines expressing genetically encoded fluorescence proteins or costly SBFSEM equipment.

**Significance statement -sentence summary:** This work describes and compares three different strategies to obtain accurate volumes of leaf plastids from Arabidopsis, the most widely used model plant. We hope our contribution will support quantitative metabolic flux modeling and spark other projects aimed at a more metric-driven plant cell biology.

## Introduction

Photosynthesis, the light-driven CO_2_ fixating pathway, is the foundation of life and global food production. In land plants, this pathway is housed in the chloroplast, a specialized plastid-type of endosymbiotic origin located in mesophyll or bundle sheath leaf cells. Chloroplasts are only one of several plastid-types found in the various diverse plant tissues (Choi et al., 2021), with the proplastid representing the most basic undifferentiated precursor organelle (Jarvis and López-Juez, 2013). Through highly coordinated gene expression involving the nuclear and the organellar genome, proplastids develop into chloroplasts, amyloplasts, chromoplasts etc., all with distinct morphologies and varying sizes (Liebers et al., 2017; Sun et al., 2017). Depending on plant age some plastid types can interconvert (Jarvis and López-Juez, 2013). The number of plastids can surpass 100 per cell in *Arabidopsis thaliana*, contributing to about a quarter of the total cell volume (Crumpton-Taylor et al., 2012; Unal et al., 2020). The abundance of genetically identical plastids occurs through binary fission facilitated by a complex contractile FtsZ ring inside the organelle and additional plastid-dividing (PD) rings that contain proteins anchored or associated with the inner and outer envelope membrane (Yoshida, 2018; Osteryoung and Pyke, 2014; Chen et al., 2018). The discovery of this intricate machinery was investigated through several loss and gain-of-function mutants of FtsZ and PD ring components. These mutants represent invaluable research tools to understanding both organelle fission and the significance of plastid abundance and size on basic physiological responses such as light stress avoidance (Dutta et al., 2017).

Plastids carry out general and highly specialized biochemical reactions, many yielding phytohormones or their respective precursors, which are critical for plant development and stress response (Bittner et al., 2022). Unsurprisingly, understanding plastid and chloroplast function has been the focus of many scientists interested in a wide range of topics, from photosynthesis to the importance of plastids for plant environmental interactions (Kleine et al., 2021). In recent years, computational modeling of energy/metabolic flux has given new insights into the complex inner workings of the organelle (Krantz et al., 2021; Fürtauer et al., 2018). This modeling has been added by non-aqueous fractionation to determine how these organelles interact (Klie et al., 2011; Fürtauer et al., 2016; Höhner et al., 2021). Nevertheless, the efficacy of these models would be further improved with precise determination of organellar volumes. This is especially important for *A. thaliana*, arguably the most studied plant worldwide and the primary model plant system for elucidating the molecular, structural, and biochemical control of energy/metabolic fluxes (Woodward and Bartel, 2018).

Chloroplast dimensions and volumes are most often inferred using two-dimensional (2D) imaging techniques (Kunz et al., 2014; Aranda-Sicilia et al., 2016; Unal et al., 2020). Transmission electron microscopy (TEM) has been the primary method for obtaining these 2D images. However, TEM requires fixation which can result in tissue, cellular, and organellar shrinkage. For instance, spinach chloroplasts loose about 30% of their volume during the fixation procedure (Winter et al., 1994). Also, TEM imaging is error-prone since optimal imaging quality requires 60-80 nm thick sections and it is impossible to know what plane of the chloroplast is visible or the angle of the section. This means it is unclear if a given chloroplast image is a glancing section or cuts through the center, making accurate volume calculations challenging with a bias towards underestimating volumes. This uncertainty leads to considerable variation in estimated chloroplast volumes and requires large time-consuming datasets to approximate accurate chloroplast volumes even within a single cell. A recent study using wheat and chickpea demonstrated that the 2D approach of estimating chloroplast volumes is inaccurate and prone to volume underestimations (Harwood et al., 2019).

The recent application of technologies to create three-dimensional (3D) representations of leaf anatomy, including the serial block-face scanning electron microscopy (SBFSEM), has introduced alternative ways to address ongoing uncertainty in chloroplast volumes (Denk and Horstmann, 2004). In short, SBFSEM employs automated collection of serial surface images from a resin-embedded sample block. This occurs via an internal ultramicrotome that cuts a 40-80 nm thin section. The newly exposed surface of the truncated sample block is scanned to generate the next SEM image. However, the SBFSEM technology requires significant specialized instrumentation, and similar with TEM has potential difficulties with sample fixation and preparation that may result in inferior images and data misinterpretation.

Live imaging of leaf tissue using confocal microscopy can avoid the errors associated with fixation, such as chloroplast shrinkage. Confocal microscopy does, however, present its own challenges and limitations. While there is no risk of chloroplast deformation due to fixation, the relatively long wavelength of light drastically lowers the achievable imaging resolution compared to electron microscopy. Additionally, since most confocal microscopes utilize a pinhole to image optical sections inside the sample tissue, the fluorescence is scattered by the tissue it passes through before reaching the objective, lowering resolution and limiting the depth accurate imaging can be done to. As long as this is taken into consideration, however, confocal can be powerful tool as it also allows for colocalizing several fluorophores within the sample, allowing for easy visualization of one or more structures of interest.

In this study, we used different 3D imaging techniques to measure leaf chloroplast volumes in *A. thaliana* (Figure 1A). Two protocols employ confocal microscopy z-stacks using chlorophyll fluorescence or a chloroplast envelope marker, respectively, as easy to replicate, more accessible methods. The third approach is based on SBFSEM and 3D reconstruction. To validate our assays, we used three different *A. thaliana* genotypes: 1) Col-0 as a wild-type control, 2) *35s-PDV1 35s-PDV2*, which has more, smaller chloroplasts, and 3) *arc5-2*, having fewer but gigantic chloroplasts per cell (Osteryoung, 2017). The *35s-PDV1 35s-PDV2* and *arc5-2* are chloroplast division mutants and were chosen to quantify the accuracy of each volume determination approach.

**Figure 1:**
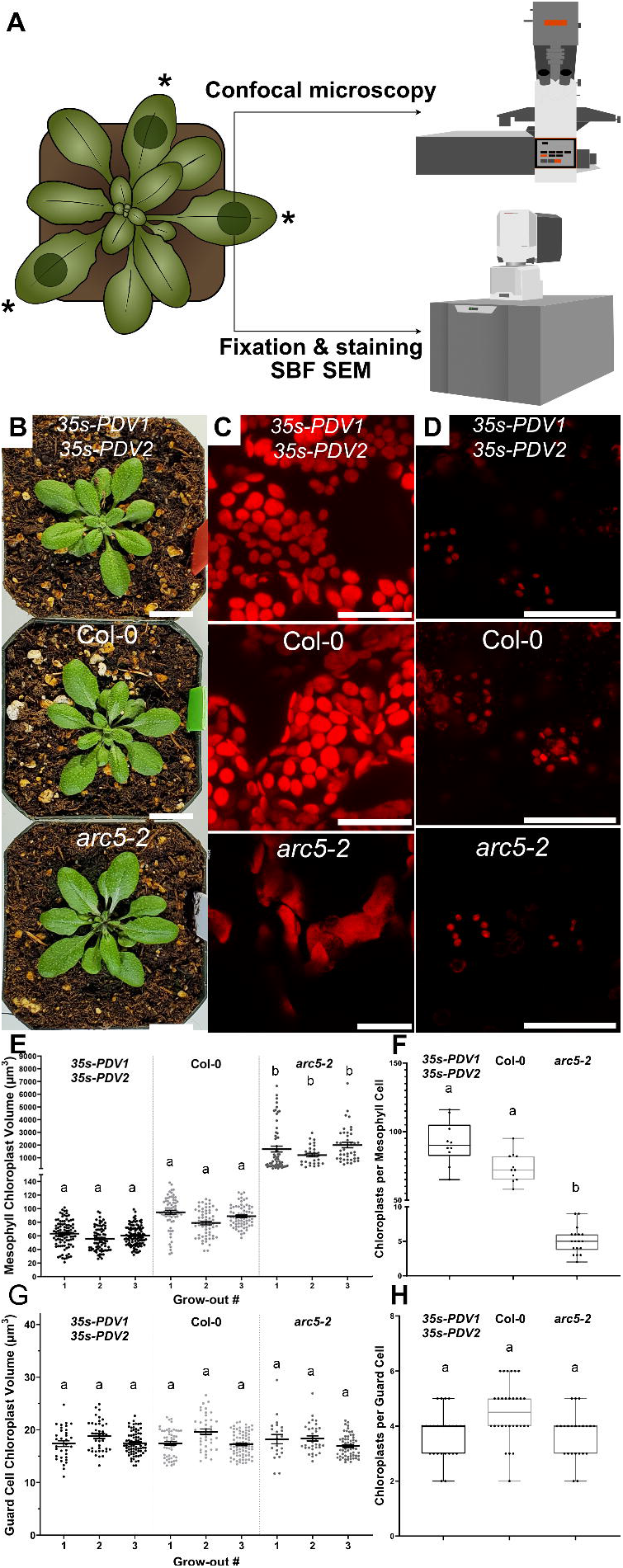
Confocal microscopy reveals changes in gain and loss of function mutants in components of the plastid division machinery that are restricted to mesophyll cells. **A,** The diagram of the Arabidopsis plant shows the size of the plants when imaging and sample preparation was done. The * indicates the leaves used for imaging, and circles indicate the location on each leaf. From here, these samples were either used immediately for imaging using confocal microscopy or fixed and prepared for SBFSEM. **B**, No apparent changes in plant growth or appearance were observed in long-day cultivated genotypes used in this study, *35s-PDV1 35s-PDV2*, Col-0, *arc5-2* (scale bars = 2 cm). **C**, Confocal images of chlorophyll fluorescence for each genotype show clear size differences in mesophyll chloroplasts (scale bars = 25 µm). **D**, Confocal images of guard cell chloroplasts taken using chlorophyll fluorescence shows similar chloroplast volumes within guard cells of the three genotypes investigated (scale bars = 25 µm). **E**, Comparison of mesophyll chloroplast volumes deduced from z-stack of chlorophyll fluorescence recordings. Individual grow-outs are shown for each genotype. Statistical analysis done by one-way ANOVA with Tukey’s multiple comparison test (*p*<0.05) showed no significant difference between Col-0 and *35s-PDV1 35s-PDV2* chloroplast volumes, neither in the full dataset, nor between any individual repetitions. Significant differences were observed between *arc5-2* and *35s-PDV1 35s-PDV2* but also between *arc5-2* and Col-0 mesophyll chloroplast volumes. Letters above each sample in the graph indicate groups of significance (±SE). **F**, Chloroplast numbers per mesophyll cell were counted and show an inverse relationship to that of chloroplast volumes observed between the genotypes. *35s-PDV1 35s-PDV2* has the highest average number of chloroplasts per cell with around 90 chloroplasts, followed by Col-0 averaging slightly fewer, at around 70 chloroplasts per cell. Finally, *arc5-2* mutants showed by far the lowest chloroplast count per cell with an average of 5 chloroplasts (±SE). **G**, Guard cell chloroplast volumes calculated based on z-stacks of chlorophyll fluorescence recordings, in three separate grow outs, statistical analysis shows no significant difference (*p*<0.05) between any of the genotypes, neither within the whole dataset nor within individual growing cycles. Letters above each data set again show groups of significance (±SE). **H**, Guard cell chloroplast counts show that, unlike the similarities in volumes, a slight difference in average chloroplast number per guard cell can be observed. Col-0 guard cells contain on average 4.5 chloroplasts, while both the *35s-PDV1 35s-PDV2* and *arc5-2* genotypes contain only around 3.5 chloroplasts per guard cell (±SE). For each data set, plastid volumes were collected from three plants per grow out (*n*=3).

## Results & Discussion

### Mesophyll chloroplast volume measurements using chlorophyll fluorescence

For plastid volume calculations three different, wild type and previously described genotypes were selected. *35s-PDV1 35s-PDV2* has smaller chloroplasts caused by the overexpression of the outer envelope PLASTID DIVISION1 (*PDV1*) and PLASTID DIVISION2 (*PDV2*) proteins, which recruit the ARC5/DRB5P ring during chloroplast division (Okazaki et al., 2009; Dutta et al., 2017). Conversely, *arc5-2* is a T-DNA insertion loss of function mutant that exhibits a low number of gigantic chloroplasts per cell. The *ARC5* locus encodes one of the outer envelope membrane proteins responsible for assembling the most outer PD ring (Robertson et al., 1996; Miyagishima et al., 2006). Col-0 was used as a wildtype control. All genotypes exhibited a similar green leaf color and were indistinguishable from controls with regards to their growth rate and appearance (Figure 1B). The reported chloroplast phenotypes became visible in the micrographs (Figure 1C-D). For all experiments, leaf discs were collected from the first three mature true leaves. Three separate grow-outs per genotype were utilized to test the consistency of our results. Initially, chloroplast volumes were calculated based on confocal microscopy z-stacks of chlorophyll fluorescence (Movie S1-3), which emerges mostly from stacked grana thylakoids (Figure 1E). Across all three grow-outs, Col-0 mesophyll chloroplasts had an average volume of 88.24 ± 1.58 µm^3^. *35s-PDV1 35s-PDV2* showed a clear trend towards slightly smaller mesophyll chloroplast volumes with an average volume of 60.13 ± 1.05 µm^3^. Lastly, *arc5-2* chloroplast volumes were significantly greater (one-way ANOVA, *p*<0.05) as in wild-type and *35s-PDV1 35s-*PDV2 plants, averaging at 1538 ± 145 µm^3^. Frequency distribution plots of the combined volume data collected on all three genotypes are shown in Figure S1A-B. Comparing different grow-outs, minor differences in chloroplast volumes can be observed within each genotype. For all three genotypes, the second grow-out season gave rise to slightly lower average chloroplast volumes than the first and third season indicating minor seasonal effects. However, statistical analysis within each genotype did not indicate significant differences between grow-outs (one-way ANOVA, *p*<0.05).

On average, Col-0 contained 74 ± 3 chloroplasts per mesophyll cell (Figure 1F). This is in the middle range compared to reports by others suggesting averages of 60, 76 (± 5) or between 80 and 120 chloroplasts per cell (Kinsman and Pyke, 1998; Okazaki et al., 2009; Crumpton-Taylor et al., 2012). In line with the literature, *35s-PDV1 35s-PDV2* and *arc5-2* mesophyll cells contained on average 92 ± 5 and 5 ± 0.5 chloroplasts, respectively (Miyagishima et al., 2006; Okazaki et al., 2009). The differences between studies can be related to different mesophyll cell types, developmental states, or to local growth conditions which were all reported to affect plastid numbers (Antal et al., 2013).

### Guard cell plastid volume measurements using chlorophyll fluorescence

Apart from the mesophyll, leaf chloroplasts can be found in vascular parenchymal cells and within the epidermal layer in pavement and guard cells (Barton et al., 2016). Guard cell chloroplasts are much smaller than mesophyll chloroplasts (Pyke and Leech, 1994). Thus, we also assayed guard cell plastid volumes to test the feasibility of our protocol across different cell types. When comparing guard cell chloroplast volumes, no differences between genotypes were observed. For Col-0 chloroplast volumes averaged at 17.69 ± 0.21 µm^3^, while volumes for *35s-PDV1 35s-PDV2*, and *arc5-2* were 18.04 ± 0.26 µm^3^, and 17.26 ± 0.28 µm^3^ respectively (Figure 1G). Comparing the genotypes by one-way ANOVA showed no statistically significant difference (*p*<0.05). Figure S2 shows an overlapping frequency distribution for the three genotypes’ guard cell plastid volumes. When comparing the three grow-outs individually the same pattern can be seen. In summary, our data are in line with previous reports on *arc5* mutants showing that loss of *ARC5* affects mesophyll but not guard plastid sizes (Pyke and Leech, 1994).

As shown above, guard cell chloroplast volumes are similar between Col-0 and the two mutants affected in mesophyll chloroplast division. Similarly, the number of plastids per guard cell was not significantly different (Figure 1H): Col-0 guard cells contained 4.5 ± 1.0 chloroplasts on average, which aligns with previous research done on this (Fujiwara et al., 2019), while both *35s-PDV1 35s-PDV2* and *arc5-2* contained 3.6 ± 0.83 slightly fewer chloroplasts on average. Some previous studies reported immature, non-fluorescing plastids in *arc5-2* guard cells (Fujiwara et al., 2018). Since no brightfield images were acquired in this study, we cannot comment on this observation as we only visualized fluorescing plastids. Also, giant chloroplasts were previously found in rare instances in *arc5-2* guard cells (Fujiwara et al., 2018), which we did not encounter.

### Mesophyll chloroplast volume measurements using a plastid outer envelope fluorescence marker protein

Chlorophyll fluorescence emerges primarily from photosystem II located in grana thylakoids and thus may not effectively represent the entire volume of the chloroplast. Therefore, a cyan-fluorescence protein (CFP (mTurquoise version)) envelope marker localized to the outer chloroplast envelope was employed. Initially, a Col-0 wild-type line with robust CFP signal was isolated using confocal microscopy (Figure 2A-C). Since strong overexpression of envelope proteins can result in membrane alterations, we ensured that this did not occur in the lines selected for this study (Breuers et al., 2012). Subsequently, the marker gene was introgressed into the *35s-PDV1 35s-PDV2* and *arc5-2* mutants with known plastid volume differences. Once homozygous F3 plants were available, we acquired z-stacks of both, the envelope CFP marker and chlorophyll fluorescence signals in parallel from all three genotypes. As expected, average chloroplast volumes calculated using the envelope marker was slightly higher than that calculated using chlorophyll fluorescence (Figure 2D). For Col-0 the average volume for chlorophyll fluorescence and the envelope marker were 78.9 µm^3^ ± 2.6 and 94.7 ± 2.9 µm^3^ respectively, for *35s-PDV1 35s-PDV2* they were 55.6 µm^3^ ± 2.4 and 63.1 ± 2.0 µm^3^ respectively, and for *arc5-2* the values were 1714.9 µm^3^ ± 222.7 and 2232.0 ± 330.0 µm^3^, respectively. Statistical analyses comparing chlorophyll fluorescence and envelope marker volumes within each genotype, did not show a significant difference in values (*p*<0.05), albeit a trend was clear. When comparing different genotypes, the same observation emerged i.e., statistically significant higher volumes exist only between *arc5-2* and the two other lines. Col-0 volumes based on chlorophyll fluorescence reflect to the lower end found in the three, independent grow-out experiments performed previously (Figure 1E). Seasonal effects are the most likely explanation.

**Figure 2:**
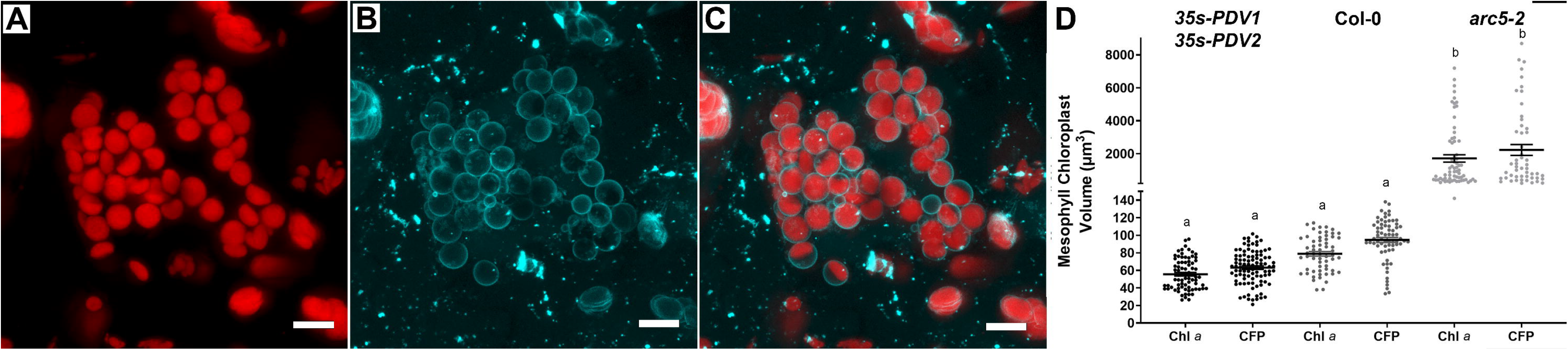
Comparing mesophyll chloroplast volumes deduced from chlorophyll fluorescence and an envelope membrane located fluorescent marker. Comparison of **A**, chlorophyll fluorescence, **B**, the outer chloroplast envelope labeled with CFP visualized in **C**, as an overlay in Col-0 (scale bar = 10 µm). **D**, comparison between chloroplast volumes extracted by chlorophyll fluorescence and CFP envelope marker shows slightly higher volumes in each genotype when calculated using the CFP marker. A statistically significant difference (*p*<0.05) in chloroplast volumes was only found between Col-0 and *35s-PDV1 35s-PDV2* when compared to *arc5-2* mutants. No significant difference is observed between imaging methods when looking at individual genotypes (±SE).

### Chloroplast volume determination using serial block-face scanning electron microscopy

When it comes to the application in plant tissues, serial block-face scanning electron microscopy (SBFSEM) is still a relatively new imaging technique. In fact, for *A. thaliana* mesophyll cell organelles there are no published volume data yet. Thus, we performed SBFSEM as an orthogonal, more high-resolution method for chloroplast analysis. Imaging was done for the same three genotypes as described above. Since the tissue has to be fixed, dehydrated and embedded, osmotic alterations may have an influence on organelle volume. The protocol used here and described in the materials and methods section was developed as part of a study to improve SBFSEM protocols and analysis in plant tissues (Mullendore et al. in preparation). This includes a machine learning algorithm “ANATOMICS MLT” which was used for this study to auto-label large image data sets. Figure 3A shows chloroplasts labeled by ANATOMICS MLT. The average chloroplast volume determined for Col-0 was 92.9 ± 1.3 µm^3^. For *35s-PDV1 35s-PDV2*, and *arc5-2* the volumes were 69.4 ± 0.8 µm^3^, and 1970 ± 759 µm^3^, respectively (Figure 3B-C). As for all previous imaging techniques used, only trends in chloroplast volumes difference could be identified between *35s-PDV1 35s-PDV2* and Col-0, while both genotypes were significantly different from *arc5-2*. A full image stack reconstruction for each genotype can be found as Movie S4-6.

**Figure 3:**
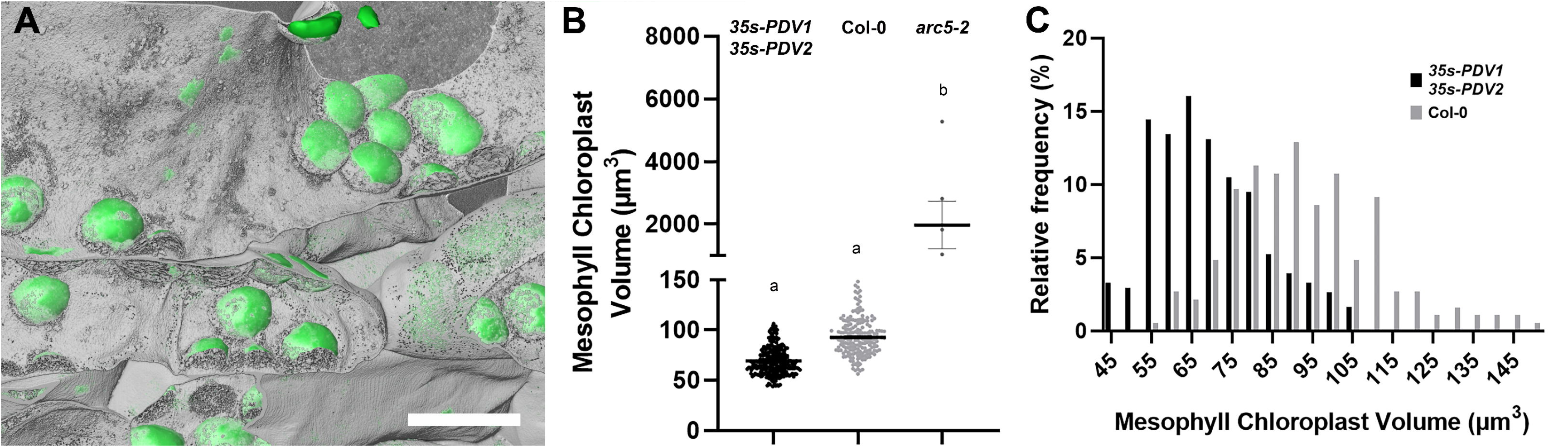
Chloroplast volume analysis using SBFSEM. **A**, Reconstructions of Col-0 chloroplasts by using SBFSEM. Green objects represent chloroplasts labeled using the machine learning algorithm, while in grey a surface was created by thresholding, representing mostly the cell wall and some other high contrast objects within the cells (scale bar = 12.5 µm). **B**, Deduced chloroplast volumes measured by SBFSEM imaging. Statistical significance (*p*<0.05) in chloroplast volumes is shown by numbers above each dataset in the graph. A significant difference in chloroplast volumes can only be seen between Col-0 and *arc5-2*, as well as between *35s-PDV1 35s-PDV2*, and *arc5-2*. **C**, Volume frequency distributions of Col-0 and *35s-PDV1 35s-PDV2* shows a shift in volume distribution between the two genotypes (±SE).

### Protocol comparisons and recommendations

In this study, we set out to gather accurate volume information from mesophyll and guard cell plastids of the model plant *A. thaliana*. In addition, we wanted to test the feasibility of employing standard confocal microscopy, available in most biology departments, to determine organellar volumes. When comparing between imaging approaches, chloroplast volumes were relatively consistent (about 93 µm^3^ in wild-type plants), with volumes derived by an outer envelope marker protein showing a slightly higher average between 10-20% when compared to both chlorophyll fluorescence and SBFSEM measurements (Figure 4A-C, Table S1). However, when running a one-way ANOVA comparing chloroplast volumes measured by each of the three imaging techniques, no statistically significant difference (*p*<0.05) was found within either of the genotypes, regardless of the imaging approach applied. Slightly higher average volumes using data from the envelope marker are to be expected, as it defines the outer boundary of a chloroplast and thus encompasses envelope membranes, inter membrane space, and the stroma. In contrast, chlorophyll fluorescence primarily represents the volume determined by thylakoid membranes. One limitation we observed when subtracting the chlorophyll fluorescence from the outer envelope-based volume is that the obtained values are too low (≈ 16 µm^3^) to accurately reflect the stromal volume. According to the literature the stroma is expected to occupy about 50% of a plastid (Antal et al., 2013). This shortcoming can be explained by the insufficient resolution of confocal micrographs which does not allow to accurately resolve the stroma between thylakoid membranes.

**Figure 4:**
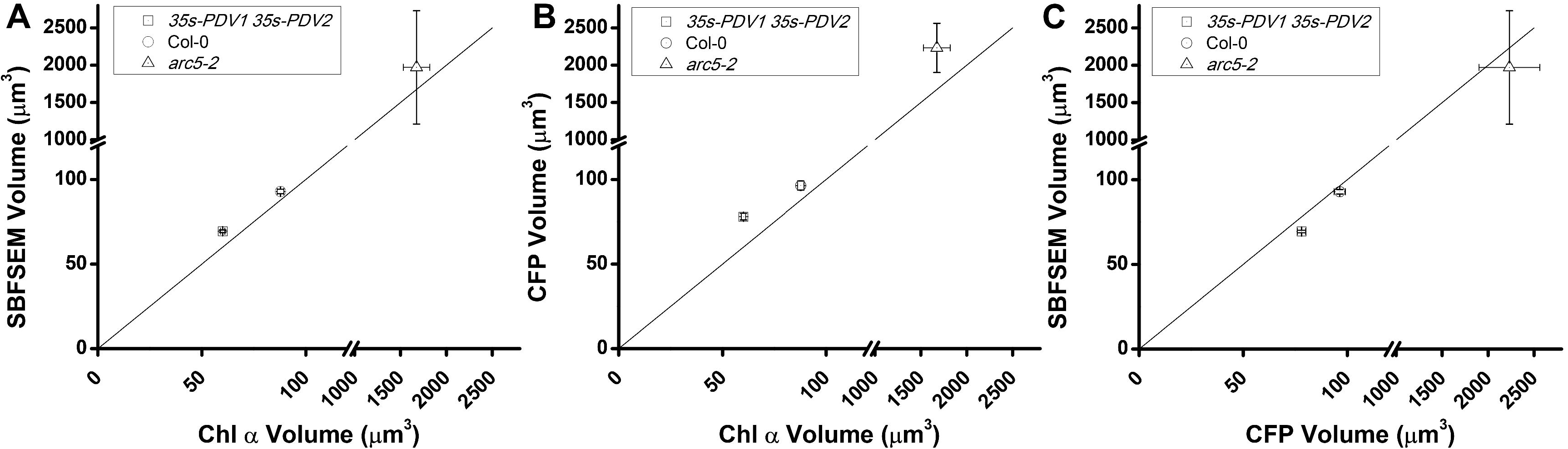
Comparison of chloroplast volumes measured by imaging type. **A-C**, comparison of mesophyll chloroplast volumes extracted using confocal imaging of chlorophyll fluorescence (Chl *a*), CFP envelope marker, and SBFSEM. The means of the data collected using each imaging method was taken for all genotypes. Each pair of imaging methods is compared by plotting the means (±SE). An identity line was added to highlight differences in means between imaging types in each graph. This shows overall similar volumes and distributions between the methods, with data extracted using the CFP envelope marker showing the largest volumes and chlorophyll fluorescence having the lowest. A one-way ANOVA done for statistical analysis only showing a significant difference (*p*<0.05) when comparing *35s-PDV1 35s-PDV2* and Col-0 volumes acquired by either method to *arc5-2*. No significant difference was found between imaging methods for any genotype.

The critical advantage of SBFSEM is the much higher resolution compared to confocal or super-resolution microscopy, but much faster acquisition time than focused ion beam methodologies. Therefore, it allows to image large tissue areas with hundreds or thousands of cells within reasonable time (days) while allowing to acquire surface areas and volumes of organelles that are too small to image by light microscopy-based methods (e.g., confocal) such as the ER or Golgi apparatus. It is however critical that appropriate protocols are used to maintain the volume of organelles. Chloroplasts are large enough to serve as a tool to compare volumes taken by confocal microscopy and SBFSEM. It is reasonable to assume that if chloroplast volumes between in situ and embedded tissue are similar, other organelle volumes in the embedded tissue should be accurate as well.

The difference in chloroplast volumes between chlorophyll fluorescence and SBFSEM images is surprisingly small and not statistically significant, showing that our protocol provides excellent maintenance of the tissue. All chloroplasts in our SBFSEM micrographs appear intact (no sharp edges which are usually an indication of shrinkage) and undamaged. Nevertheless, fixative and buffers concentrations as well as the dehydration procedure may have to be tailored to other specimen types such as stems and roots or other plant species. Stains can also be adjusted to fit organelles or structures of interest. Usually this requires a fair amount of strategic trial and error. However, SBFSEM currently allows standard resolutions of down the 10 nm at comparably high acquisition speeds which is a massive improvement over other available technology. As we show in this study, accurate quantitative anatomical data comparable to in situ studies can be achieved.

The chloroplast division mutants used as a proof of concept in this study demonstrates the ability and limitations to quantify volume differences. While each method confirmed the drastic mesophyll plastid size variation between Col-0 and *arc5-2* plants, the expectedly smaller differences between Col-0 and *35s-PDV1 35s-PDV2* were only revealed by trend but not with sufficient statical power to allow strong conclusions regardless of the imaging applied. Volume differences between mesophyll and guard cell chloroplasts were also successfully resolved. Interestingly, while mesophyll chloroplast volumes varied among genotypes, guard cell plastids had consistent volumes and numbers throughout. This provides more evidence that guard cell and mesophyll plastid division are governed by distinct genetic programs.

When comparing our results with the literature we find quite stark differences. While we determined the Col-0 wild-type mesophyll chloroplast using orthogonal approaches at a size of ≈ 93 µm^3^ most publications refer to old volume studies on other species and an average value of ≈ 31 µm^3^ (Antal et al., 2013; Nobel, 2020). Although mesophyll chloroplast sizes seem highly species- and to a minor degree daytime-dependent the values from spinach, pea, tobacco, wheat, poplar are in the range from 15 to 35 μm^3^ and appear to be rather underestimations (Antal et al., 2013; Nobel, 2020). At least side by side in a light microscope *A. thaliana* and pea chloroplast do not seem to drastically differ in size (Schulz et al., 2004). Recent 3D assays describe wheat chloroplasts at 114.6 ± 21.5, chickpea at 22.4 ± 10.2 µm^3^, and rice at 47 µm^3^ (Oi et al., 2017; Harwood et al., 2020). For *A. thaliana* only one study on cotyledon development was found to report 3D derived plastid volume data (Pipitone et al., 2021). Albeit only three plastids were assayed by SBF-SEM the obtained value of 112.14 µm^3^ (±4.3) is roughly in the same range as the 93 µm^3^ we recorded from mesophyll chloroplasts in mature true leaves. In conjunction, both studies emphasize that *A. thaliana* chloroplasts are far bigger than the often-suggested average plastid volume of 31 µm^3^. This needs to be taken into account when organelle volumes are employed in flux models. Additionally, because of the wide range of plastids sizes reported from different species generalizing metabolic flux assumptions can be problematic. Fortunately, as we show through our study rapid z-stack recordings based on chlorophyll fluorescence using a standard confocal microscope gives sufficiently accurate volume information to survey this data point for any given plant species of interest. Since we obtained very similar values regardless of the approach employed, we can state with high confidence that a volume of ≈ 93 µm^3^ reflects the natural in situ situation in the model plant Arabidopsis. Moreover, the fixation-staining protocol we utilized for SBFSEM is well-suited for plastids. The low deviation from values obtained via fixation-staining-free confocal microscopy confirms that no shrinkage occurred in our specimens.

## Conclusions

Overall, comparing chloroplast volumes from plastid division mutants acquired through different imaging methods shows relatively close overlap of volumes. This suggests that even though z-stack confocal micrographs have much lower resolution than SBFSEM, they still give fairly accurate volume data. Between the two confocal microscopy methods, using an envelope marker will yield more accurate total chloroplast volumes. Nevertheless, for a much easier and faster estimation, chlorophyll fluorescence delivers high replicate numbers and is thus quite accurate. Moreover, it does not require cloning, transformation or introgression of a fluorescence protein encoding transgene. One caveat is that chlorophyll fluorescence-based volume determination is not recommended for mutants with affected chlorophyll metabolism. SBFSEM yields the most accurate volume data due to its much higher resolution. However, lengthy sample preparation procedure optimization might be necessary to prevent artifacts resulting in skewed volumes. Since plastid volumes depend on light conditions, growth temperatures, and the genetic makeup of plants we encourage more research on this subject using the simple protocols introduced here. Scientists are advised to consider our conclusions for balanced and informed decision-making which answer the question at hand with the best equipment available to them.

## Experimental procedures

### Plant Material

*A. thaliana* plants used: Columbia-0 as a control, *arc5-2* (SAIL_71_D11), a giant chloroplast mutant due to a knockout of the *ARC5* chloroplast division gene, and 35*S-PDV1 35S-PDV2* with smaller chloroplasts caused by overexpression of *PDV1* and *PDV2* (Miyagishima et al., 2006; Okazaki et al., 2009; Dutta et al., 2017). These plants were grown in a Conviron growth chamber (Winnipeg, Man, Canada) using a 16/8-hour light/dark cycle at 150 µmol photons m^-2^s^-1^, 21°C/19°C day/night cycle and 60-80% humidity on soil. Approximately, 100 seeds per genotype were sown into one 1.15-quart pot with soil (Sungro Professional Growing Mix #1, Sun Gro Horticulture, Agawam, MA, USA), and after a week, the ten largest seedlings were transferred into individual 0.59-pint pots and grown for 4 additional weeks before imaging.

To target a fluorescent marker to the outer chloroplast envelope surface, the coding sequence (192 bp) of the outer envelope protein 7.1 (OEP7; At3g52420) without its Stop codon and with a Gly-Gly-Ser-Gly-linker at the 3’-end was first amplified using Q5 High-Fidelity DNA Polymerase (New England Biolabs) and the primers GGB_OEP7_F – AACAGGTCTCAAACAATGGGAAAAACTTCGGGAGC and GGC_OEP7_GGSG_R – AACAGGTCTCTAGCCTCCAGATCCTCCCAAACCCTCTTTGGATGTGG, followed by FastDigest Eco31I (Thermo Fisher Scientific) digestion, and ligation into the GreenGate module backbone pGGB (Lamproupolos et al., 2013). Together with previously described GreenGate modules (Lamproupolos et al., 2013; Waadt et al., 2017), OEP7, the orange fluorescing mNectarine (Johnson et al., 2009) without Start codon, and the cyan fluorescing mTurquoise (Goedhart et al., 2010) with a 5’-end Gly-Ser-linker were ligated into the plant expression vector pGGZ003 yielding the plasmid pGGZ-RW105 (pGGZ003-pUBQ10-OEP7-GGSG-mNectarine_ATG-GSL-mTurquoise-tHSP18.2M-hygR). This construct was then transformed into Col-0 wild-type by floral dip, and positive transformants were selected through germination on hygromycin (15 μg ml^-1^, Chem-Impex, Wood Dale, IL, USA). Subsequently, the envelope marker was introgressed into *arc5-2* and *35s-PDV1 35s-PDV2* mutant lines.

## Confocal Laser Scanning Microscopy

### Mesophyll Chloroplast Fluorescence Volume Measurements

For mesophyll imaging, fresh leaf disks of 1 cm diameter were taken from the center of a mature leaf and imaged from the lower epidermal side with a HC PL APO CS2 63x/1.20 NA water-immersion objective on a Leica SP8 confocal laser-scanning microscope (Leica Biosystems, Deer Park, IL, USA). Chlorophyll was excited using a 405 nm pulsed laser and the fluorescence was collected using a HyD detector set to 650-720 nm. The gain was optimized for each stack, using the brightness indicator. Additional imaging parameters were: scanning speed 100 Hz, zoom 2, line average 2, pinhole 1, and z-stacks system optimized with 0.305-µm-thick optical sections for volume analysis. Number of z-sections varied, and stacks were later cropped to exclude images where the fluorescence border could not be accurately identified.

### Guard Cell Chloroplast Volume Measurements

For imaging of guard cell chloroplasts, the abaxial side of the leaf was used as there are more stomata present. The abaxial side of a leaf was glued to a glass slide using medical adhesive (Hollister, Libertyville, IL, USA). After allowing the glue to dry, leaf tissue was removed by scraping with a razor blade until the lower epidermis became exposed (Azoulay-Shemer et al., 2016). Guard cells turgescence was determined visually and used to checked for intactness, and guard cell chloroplasts were then imaged using a HC PL APO CS2 63x/1.40 oil-immersion objective. Z-stacks were taken using 0.255-µm-thick optical sections. Otherwise, images were taken as described above.

### Cyan-fluorescence Protein Imaging

Plants with chloroplast envelopes tagged with CFP (mTurquoise) were also imaged using a HC PL APO CS2 63x/1.20 NA water-immersion objective on a Leica SP8 confocal laser-scanning microscope (Leica Biosystems, Deer Park, IL, US). Dual channels were used to image CFP simultaneously with chlorophyll fluorescence. A HyD detector set to 460-520 was used to collect CFP fluorescence, while a HyD detector set to 650-720 nm was used for chlorophyll fluorescence. All other imaging parameters were as described above.

### Chloroplast Volume Analysis

Chloroplast reconstruction and analysis was done using ImageJ software, utilizing the Bio-Formats plugin to import .lif files (https://github.com/ome/bioformats). After importing a z-stack, and manually thresholding to optimize chloroplast outlines, a Median Filter (Radius: 2.0) was applied for smoothing and subtract (∼10), a global brightness reduction tool, was used to remove background noise. Chloroplast volumes were extracted with the 3D Objects Counter function in ImageJ, and the resulting labels were checked to ensure only volumes of complete chloroplasts were counted.

### Mesophyll Cell Chloroplast Count

To count the chloroplasts per cell in mesophyll cells, z-stacks were taken using a 20x water-immersion objective. Two channels were used to image chlorophyll autofluorescence and brightfield simultaneously, allowing us to distinguish the borders of cells. Z-stacks were filtered as detailed above, reconstructed, and chloroplasts counted.

## Serial Block Face Scanning Electron Microscopy (SBFSEM)

### Fixation Protocol

Arabidopsis leaves were removed from the plant, cut into 2 mm x 2 mm squares, and put into a fixative solution containing 4% glutaraldehyde, 2 mM CaCl_2_ in 0.1 M cacodylate buffer (pH 6.8) for 6 hours at room temperature. Samples were then microwaved at 300W at a 35°C maximum temperature limit for 2 min and then washed 3x for 10 minutes in 0.1 M cacodylate buffer followed by a post-fixation with 1.5% K_4_Fe(CN)_6_, 2% OsO_4_, 2 mM CaCl_2_, in 0.15 M cacodylate buffer overnight at 4°C. After washing 3x for 10 min in double distilled water (ddH_2_O) at room temperature, samples were incubated in 0.2% gallic acid for 1 hour at room temperature, and then washed 3x for 10 minutes in ddH_2_O. A secondary post-fixation was done in 2% OsO_4_ for 3 hours at room temperature, and after washing 3x for 10 min in ddH_2_O, a 2% uranyl acetate incubation was applied overnight at 4°C. Samples were washed 3x for 10 minutes in ddH_2_O at room temperature, then stained with Walton’s lead aspartate (Walton, 1979) at 60°C for one hour and finally washed 3 times for 10 min in ddH_2_0 at room temperature.

Samples were dehydrated with an acetone series using freshly mixed acetone solutions. 10% steps were done between 10%-50% (v/v) acetone for 10 min at room temperature. A second exchange of 50% acetone was incubated at -20°C for 1 hour. For the remainder of the dehydration series, exchanges were performed at -20°C overnight in 60%, 70%, 80%, 90%, 100%, 100%, 100% (v/v) acetone. After the third 100% acetone treatment, samples were incubated overnight and then moved to room temperature to acclimate for ∼30 min before the final dehydration in 100% acetone two times for 10 minutes at room temperature.

Samples were infiltrated in Spurr’s resin (Sigma Aldrich, St. Louis, MO, USA) in 3:1, 2:1, 1:1, 1:2, 1:3, 100% acetone:hard Spurr resin without the hardener of the resin (DMAE) to prevent premature polymerization, each overnight at room temperature on a rotator. This was followed by two exchanges of 100% hard Spurr’s resin with DMAE, overnight at room temperature on a rotator, with the lids removed. Finally, samples were microwaved at 100W and 40°C for 1 hour and subsequently put into a 70°C oven to polymerize.

### Imaging

To prepare samples for SBFSEM, the embedded tissue was trimmed and sectioned using a Leica EM UC7 ultramicrotome. Ultrathin sections (∼70 nm) were taken and checked for quality using a FEI Technai G2 20 Twin (Thermo Fisher, Waltham, MA, USA). Samples were then transferred to a SBFSEM stub, which is the sample holder, trimmed on the ultramicrotome using a glass knife, after which a Technics Hummer V Sputter Coater was used to apply 10 nm of gold coating. For imaging, an Apreo VolumeScope SEM (Thermo Fisher, Waltham, MA,USA) with the VS-DBS: LoVac lens-mounted BSED detector was used. Imaging conditions were set to 2 kV accelerating voltage, 50 Pa chamber pressure, a beam current of 0.10 nA, a pixel size of 20 x 20 nm, and a dwell time of 3 µs.

### Image Processing

The SBFSEM image stacks were processed for volume analysis using software specifically designed to work with SBFSEM stacks, Amira (Thermo Fisher, Waltham, MA, USA). Partial stacks (∼500 images) were imported into Amira, and a gaussian filter (XY, standard deviation 1, 1, kernel size factor 2) was used to remove noise. Images are then inverted, resampled (Lanczos Filter, Voxel Size: x= 40, y= 40, z= 40), and then exported as a sequence of 2D tiff files. This process was repeated until the entire SBFSEM stack had been processed and exported as described above. The complete stack of processed images was then imported into Amira, cropped around the area of interest and contrast matched (XY planes, mean & variance) using the best contrasted image within the stack as a reference. This made the contrast consistent for all images throughout the SBFSEM stack. Auto align slices (Rigid, Align and Resample) was run to automatically align all SBFSEM slices. Images were cropped again to remove overlapping edges created by the alignment, and a median filter (XY planes, 3 iterations, Iterative) was used to remove remaining noise. Finally, the stack was exported as a set of 2D tiff files. The fully processed stack was then imported into ImageJ as an image sequence, manually contrasted, and saved as a tiff, creating a 3D tiff stack.

### Volume Analysis

Volume analysis of SBFSEM stacks was done using Anatomics MLT (https://github.com/ajbrookhouse/WSU_PlantBio_ML), a machine learning algorithm designed to analyze volumes, object counts, and surface areas of features within SBFSEM stacks. Amira was used to label all chloroplasts within a sequence of 10 processed images with the Segmentation tool by using the brush, then images and labels were exported as separate 3D tiff files. These stacks were then used in the train function of Anatomics MLT (Training Config: Instance.yaml, 100,000 iterations) to train the program to label chloroplasts. The model was then used in the Auto-Label section to label full image stacks. Finally, the Output Tools tab was designed to analyze data from the created labels and can be used to create mesh or point clouds to visually display the labels. This was used to calculate chloroplast volumes and create Point Clouds of the labels which were viewed and overlayed with original images to check for labeling accuracy using the Visualize tab.

### Statistical Analysis

Graphing and statistical analysis was done using GraphPad software (Dotmatics, Boston, MA, USA). Means as well as standard deviation and error were calculated. One-way ANOVAs were used to determine statistical significance between biological replicates, genotypes, and imaging techniques. A 2-way ANOVA was run to analyze volume data within the chloroplast fluorescence dataset, comparing each growth cycle within a genotype to the volumes of chloroplasts between genotypes. A 3-way ANOVA was also used to compare chloroplast volumes of genotypes within each imaging method and to compare volumes of chloroplasts of each genotype between imaging techniques.

### Accession numbers

*arc5-2* (SAIL_71_D11, S870785), *35s-PDV1 35s-PDV2, ARC5* (AT3G19720), *PDV1* (AT5G53280), *PDV2* (AT2G16070), *OEP7.1* (At3g52420)

## Acknowledgements

We thank Dr. Katherine W Osteryoung from Michigan State University for providing the original mutant germplasms used in this project. J.K. received the WSU Franceschi training grant to support the microscopy work. J.K. thanks his graduate committee members Drs. Andrew McCubbin and Helmut Kirchhoff, and greenhouse manager Chuck Cody (all at WSU) for their support and insights. Lastly, we thank Susanne Mühlbauer (LMU Munich) for designing the illustrations shown in Figure 1 and the graphical abstract.

## Short legends for Supporting Information

**Figure S1: A**, Frequency distribution of all mesophyll chloroplast volumes calculated based on chlorophyll fluorescence shows a distribution towards lower volumes in *35s-PDV1 35s-PDV2* than in Col-0. **B**, Frequency distribution for chlorophyll-based chloroplast volumes in *arc5-2* shows volumes below 1000 µm^3^ are the most common. However, values reach as high as 7000 µm^3^.

**Figure S2:** Frequency distribution of guard cell chloroplast volumes shows very similar distribution and overall volumes between all three genotypes.

**Table S1:** Summary of chloroplast volumes in µm3 calculated using image stacks from each method employed. For each genotype the average and standard error are presented.

**Movie S1-3:** Video reconstructions showing confocal Z-stacks of each genotype, *35s-PDV1 35s-PDV2*, Col-0 and *arc5-2*, respectively. Scale bars denote 20 µm.

**Movie S4:** Video reconstruction of scanning through a *35s-PDV1 35s-PDV2* SBFSEM image stack.

**Movie S5:** Video reconstruction of scanning through a Col-0 SBFSEM image stack. Chloroplasts are labeled using the machine learning algorithm and displayed in cyan.

**Movie S6:** Video reconstruction of scanning through an *arc5-2* SBFSEM image stack.

