## Supplemental Figures for "Probing the *in-situ* volumes of Arabidopsis leaf plastids using 3D confocal and scanning electron microscopy"

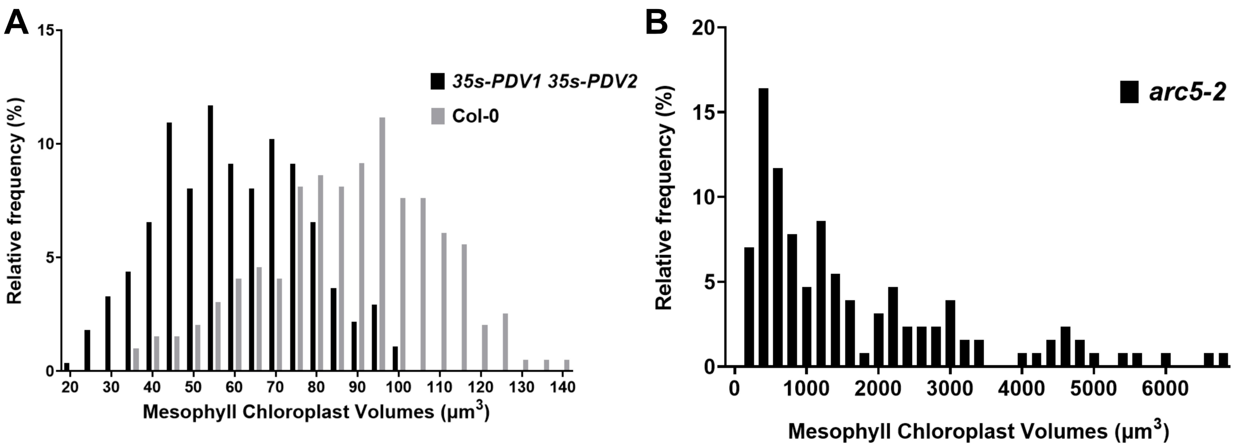


**Figure S1: A**, Frequency distribution of all mesophyll chloroplast volumes calculated based on chlorophyll fluorescence shows a distribution towards lower volumes in *35s-PDV1 35s-PDV2* than in Col-0. **B**, Frequency distribution for chlorophyll-based chloroplast volumes in *arc5-2* shows volumes below 1000 $\mu m^3$ are the most common. However, values reach as high as 7000 $\mu m^3$.

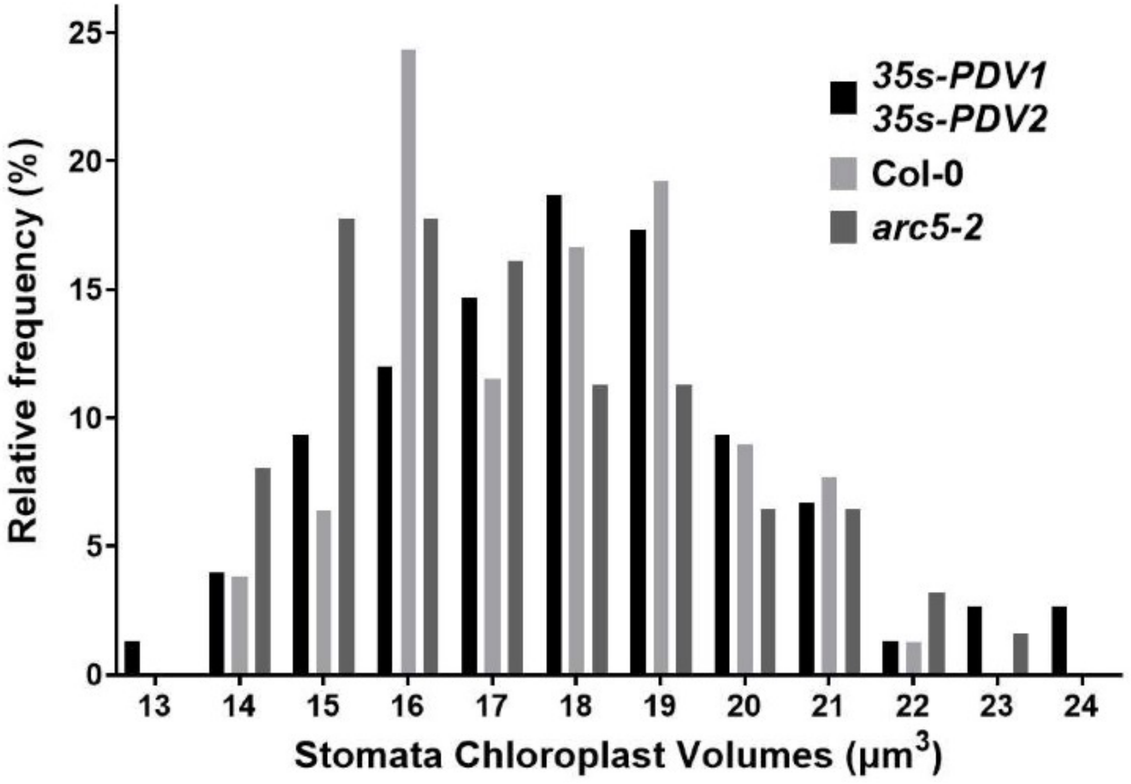


**Figure S2:** Frequency distribution of guard cell chloroplast volumes shows very similar distribution and overall volumes between all three genotypes.

**Table S1:** Summary of chloroplast volumes in μm$^3$ calculated using image stacks from each method employed. For each genotype the average and standard error are presented.

| | Average plastid volume [μm$^3$] | | |
|---|---|---|---|
| | ***35s-PDV1 35s-PDV2*** | **Col-0** | ***arc5-2*** |
| Chl a (± SE) | 60.13 (±1.05) | 88.24 (±1.58) | 1538 (±145) |
| SBFSEM (± SE) | 69.42 (±0.77) | 92.93 (±1.32) | 1971 (±759.2) |
| CFP (± SE) | 63.08 (±1.98) | 94.66 (±2.88) | 2232 (±329.7) |
| Chl a Guard Cell (± SE) | 18.04 (±0.26) | 17.69 (±0.21) | 17.26 (±0.28) |
